# Development of neonatal brain functional centrality and alterations associated with preterm birth

**DOI:** 10.1101/2022.06.01.494304

**Authors:** Sunniva Fenn-Moltu, Sean P Fitzgibbon, Judit Ciarrusta, Michael Eyre, Lucilio Cordero-Grande, Andrew Chew, Shona Falconer, Oliver Gale-Grant, Nicholas Harper, Ralica Dimitrova, Katy Vecchiato, Daphna Fenchel, Ayesha Javed, Megan Earl, Anthony N Price, Emer Hughes, Eugene P Duff, Jonathan O’Muircheartaigh, Chiara Nosarti, Tomoki Arichi, Daniel Rueckert, Serena Counsell, Joseph V Hajnal, A David Edwards, Grainne McAlonan, Dafnis Batalle

## Abstract

Formation of the functional connectome in early life underpins future learning and behaviour. However, our understanding of how the functional organisation of brain regions into interconnected hubs (centrality) matures in the early postnatal period is limited, especially in response to factors associated with adverse neurodevelopmental outcomes such as preterm birth. We characterised voxel-wise functional centrality (weighted degree) in 366 neonates from the Developing Human Connectome Project. We tested the hypothesis that functional centrality matures with age at scan in term-born babies and is disrupted by preterm birth. Finally, we asked whether neonatal functional centrality predicts general neurodevelopmental outcomes at 18 months. We report an age-related increase in functional centrality predominantly within visual regions and decrease within motor and auditory regions in term-born infants. Preterm-born infants scanned at term equivalent age had higher functional centrality predominantly within visual regions and lower measures in motor regions. Functional centrality was not related to outcome at 18 months old. Thus, preterm birth appears to affect functional centrality in regions undergoing substantial development during the perinatal period. Our work raises the question of whether these alterations are adaptive or disruptive, and whether they predict neurodevelopmental characteristics that are more subtle or emerge later in life.

## Introduction

The brain is continuously changing across the lifespan, but is most plastic and modifiable during early life (De Asis-Cruz et al. 2015). Following establishment of its general architecture during the first two trimesters of gestation, the third trimester sees vast axonal growth and rapid expansion of the cortex, the start of myelination, synaptogenesis and dendritic formation, which continues into the postnatal period (see review by Keunen et al., 2017). Alongside the establishment of the brain’s structural architecture, the formation and differentiation of functional circuits is also taking place (Cao et al. 2017), establishing a pattern of functional connectivity characterised by both segregation and integration. With such complex processes taking place during late gestation, the brain may be particularly vulnerable to factors disrupting typical maturational trajectories in this timeframe, such as preterm birth.

Preterm birth (<37 weeks’ gestation) most commonly happens during the third trimester and is estimated to affect around 11% of all live births around the world (Chawanpaiboon et al. 2019). Although improvements in perinatal care have increased survival rates, preterm infants still show higher incidence of neurodevelopmental conditions such as autism and ADHD (Wang et al. 2017; Bröring et al. 2018) and related neurodevelopmental complications in later childhood such as language delay and altered socio-emotional and behavioural functioning (Boardman and Counsell 2020). Such difficulties further translate into a significant impact on quality of life indicators such as academic achievement and peer relationships (Rogers et al. 2018). Therefore, studying both typical and atypical functional connectivity during the perinatal period may help us identify potential early biological underpinnings of developmental and behavioural phenotypes observed in preterm-born children as they grow.

Graph-theory based network methods can be applied to resting-state functional magnetic resonance imaging (rs-fMRI). This allows calculation of specific network measures such as voxel-wise degree centrality (DC). DC maps the functional connectome architecture at the voxel level by measuring the number of direct connections between a given voxel and the rest of the brain (De Asis-Cruz et al. 2015; Gozdas et al. 2018). This provides insight into the patterns of functional connectivity within the whole brain network (Zuo et al. 2012). This approach does not require *a priori* definition of regions of interest, and does not require a parcellation step, which has proven challenging in neonatal imaging (Oishi et al. 2019). As the DC of a voxel reflects the relative importance of that voxel within the network, it allows us to identify regions which are ‘central’ (“functional hubs”), and thus of likely functional importance. DC has been shown to have relatively high test-retest reliability in adults (Zuo and Xing 2014), and has offered important insight into the complexity of the adult functional connectome (Zuo et al. 2012). The measure has been employed both in the study of typical function (van den Heuvel and Sporns 2013) and it has identified consistent changes in network connectivity associated with neurodevelopmental conditions such as autism (Holiga et al. 2019), for which preterm birth is a known risk factor (Crump et al. 2021). However, studies of already diagnosed individuals, even in later childhood, cannot untangle the effect of early exposure and the secondary and/or compensatory effects of living with neurodevelopmental difficulties on the functional connectome. Studies in the earlier postnatal period are needed to understand how exposures in this period shape the developing brain.

Initial work in infants has shown that the neonatal functional connectome shows a heavy-tailed degree distribution (De Asis-Cruz et al. 2015; Oldham and Fornito 2019), with the most highly connected, and thus putative “hub” regions, located within motor, sensory, auditory and the visual cortex (Fransson et al. 2011). Efforts have also been made to understand how functional DC changes throughout the perinatal period, and how it is impacted by preterm birth. For example, neonates scanned shortly after birth between 31.3 to 41.7 weeks postmenstrual age (PMA) (Cao et al. 2017) showed age-related increases in DC predominantly within the precentral and postcentral gyri and supplementary motor area, and decreases within the posterior cingulate and precuneus cortex. However, preterm-born infants were not compared to term-born infants scanned at an equivalent age, therefore the results capture the effect of gestational age rather than the impact of pre-term birth. Only one small study has examined preterm (≤31 weeks’ gestation) and term neonates with both groups scanned at term-equivalent age. The authors reported that compared to full-term born infants, very preterm infants had higher DC in auditory and language networks compared to term infants, whereas term infants had higher centrality in frontal networks compared to very preterm infants (Gozdas et al. 2018). Moreover, the functional connectome was not examined at voxel-level resolution, which may be more sensitive to subtle alterations caused by preterm birth. Finally, no study has investigated whether functional centrality in the perinatal period is altered by preterm birth and whether this has developmental consequences later in infancy.

The Developing Human Connectome Project (dHCP) represents the largest open-source sample of resting-state fMRI neonatal data with high spatial and temporal resolution (Edwards et al. 2022), thus providing a unique opportunity to characterise functional centrality during this early phase of life. In this study, we leverage dHCP data to characterize functional centrality in the neonatal brain at the voxel-level in term-born and preterm-born neonates. We subsequently explore the relationship between neonatal functional centrality and behavioural outcome measures at 18 months of age. By understanding how functional connectivity patterns typically develop with age (postmenstrual age at scan) and how typical development is affected by the degree of preterm birth (gestational age at birth (GA)), we can both establish a typical maturation profile and investigate the consequence of preterm birth. Finally, we assessed whether functional centrality in early life is predictive of general neurodevelopmental outcomes in toddlerhood in term and preterm-born children.

## Methods

### Participants

Research participants were recruited as part of the Developing Human Connectome Project (dHCP, http://www.developingconnectome.org/). Ethical approval was given by the UK National Research Ethics Authority (14/LO/1169), and written consent was obtained from all participating families prior to data collection. For the purpose of this study, we included subjects without visible abnormality on MRI of possible/likely clinical significance following reporting by an experienced Neonatal Neuroradiologist. Findings leading to exclusion were major lesions within white matter, cortex, cerebellum and/or basal ganglia, small head/brain < 1^st^ centile, and isolated non-brain anomalies with possible clinical significance. Incidental findings with unlikely significance for clinical outcome were not excluded, such as subdural haemorrhages, isolated subependymal cysts, and punctate white matter lesions. In the case of a multiple pregnancy, only one sibling was included in subsequent analysis. Subjects were also excluded due to excessive in-scanner motion (more than 10% framewise displacement (FD) motion outliers (FD>1.5IQR+75^th^ centile)). The resulting population was 366 neonates. Of these, 300 were born at term and 66 were born preterm, i.e. before 37 weeks’ gestation. Scans were acquired at term or term equivalent age, respectively. See demographic and clinical details in Table 1. For analyses assessing the effect of PMA at scan on DC, only term-born infants were included (n=300). The effect of preterm birth was assessed by treating GA at birth as a binary variable, with comparisons carried out between the neonates born <37 weeks (preterm, n=66), and those born at >37 weeks (term, n=300). As the <37 week cut-off for preterm-birth is somewhat arbitrary (Kramer et al. 2012; Vogel et al. 2018), in addition to the dichotomous analysis, term and preterm groups were also combined (n=366), and GA at birth treated as a continuous variable.

**Table 1:**
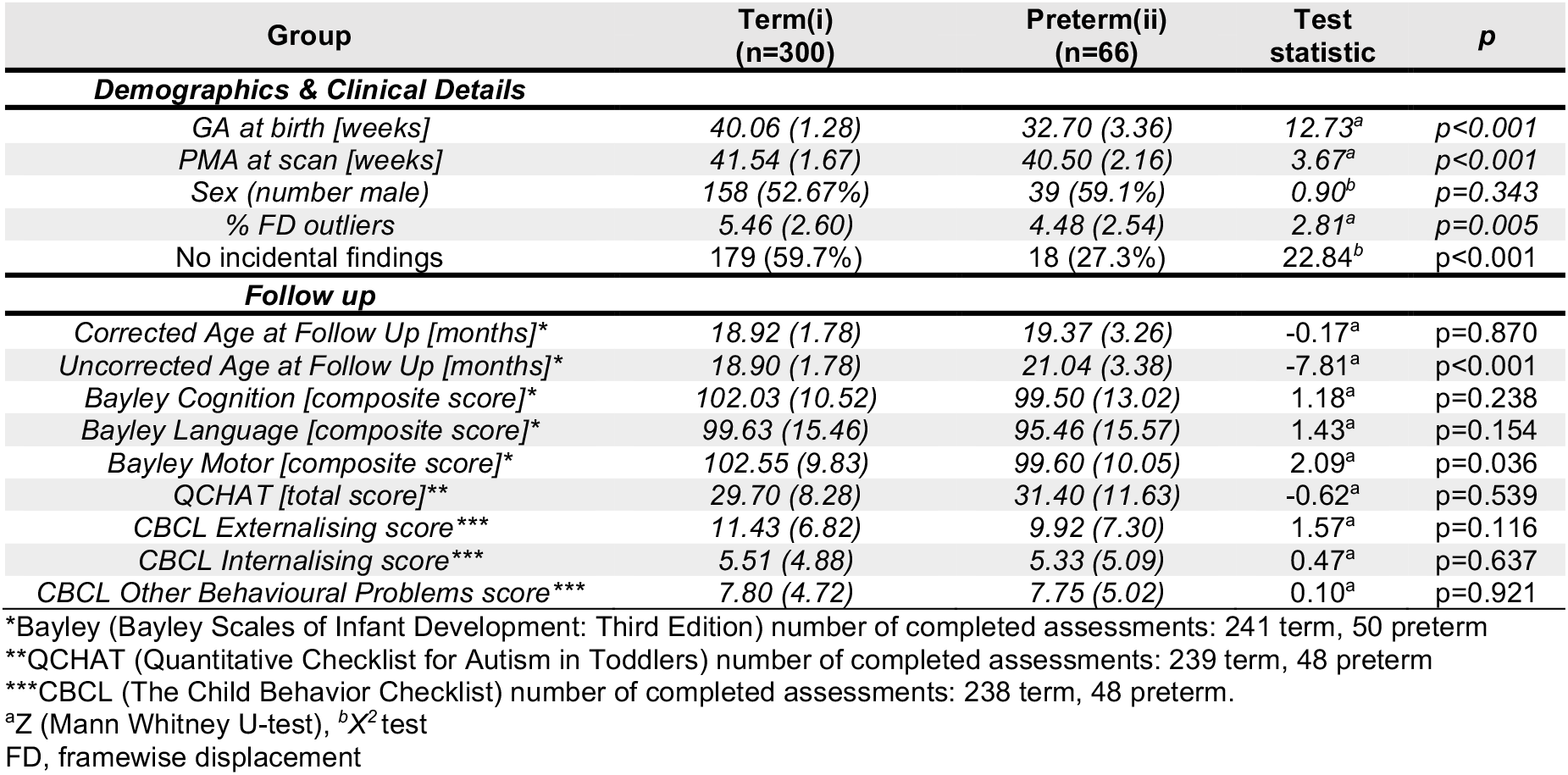
Subject Demographics, Clinical Details and Follow Up. Continuous variables expressed as mean and standard deviation. Categorical variables expressed as number and percentage.

### Image Acquisition

Infants were scanned at the Evelina Newborn Imaging Centre, Evelina London Children’s Hospital using a 3 Tesla Philips Achieva system (Best, NL) and a neonatal 32 channel receive head coil and imaging system (Rapid Biomedical GmBH, DE) (Hughes et al. 2017). Images were acquired during natural sleep following feeding, swaddling and immobilization in a vacuum evacuated mattress (Med-Vac, CFI Medical Solutions, Fenton, MI, USA). Hearing protection (moulded dental putty (President Putty, Coltene Whaledent, Mahwah, NJ, USA)) in addition to adhesive earmuffs (MiniMuffs, Natus Medical, Inc., San Carlos, CA, USA) and physiological monitoring devices (oxygen saturations, heart rate, and axillary temperature) were applied before image acquisition, and all scans were attended by a neonatal nurse and/or pediatrician. BOLD fMRI data were acquired using a multislice gradient-echo echo planar imaging sequence with multiband excitation (multiband factor 9) with the following parameters: repetition time, 392 ms; echo time, 38 ms; spatial resolution, 2.15 mm isotropic; flip angle, 34°; 45 slices; total time, 15 min 3 sec (2300 volumes). High-resolution anatomical T1-weighted images (reconstructed spatial resolution, 0.8 mm isotropic; field of view, 145 × 122 × 100 mm^3^; and repetition time, 4795 ms) and T2-weighted images (reconstructed spatial resolution, 0.8 mm isotropic; field of view, 145 × 145 × 108 mm; repetition time, 12 s and echo time, 156 ms) were acquired for registration and clinical reporting (Edwards et al. 2022).

### Functional data pre-processing

The data was pre-processed using a pipeline optimized for neonatal imaging as part of the dHCP (Fitzgibbon et al. 2020). In brief, this included dynamic distortion and intra- and inter-volume motion correction, using a pipeline including slice-to-volume and rigid-body registration. Motion, cardiorespiratory and multiband acquisition artefacts were regressed out together with single-subject ICA noise components identified using FSL FIX (Griffanti et al. 2014; Salimi-Khorshidi et al. 2014). Data was subsequently registered into T2w native space using boundary-based registration and non-linearly transformed to a 40-week neonatal template from the extended dHCP volumetric atlas (Schuh et al. 2018). Participants with high framewise displacement (FD>1.5IQR + 75^th^ centile) in more than 230 (10%) of the 2300 volumes were excluded from analysis. The data was subsequently smoothed with a 3mm Gaussian smoothing kernel, bandpass filtered at 0.01-0.1Hz and masked with a grey matter mask defined in template space.

### Functional Data Analysis

Graphs were constructed from the data at a voxel level. The BOLD time series for each voxel within the predefined grey matter mask was extracted, and a correlation matrix made using the temporal Pearson’s correlation of time series between each pair of voxels. A threshold of 0.20 was used to avoid including spurious connections in the correlation matrix (Buckner et al. 2009; Holiga et al. 2019) and only positive connections were assessed (Murphy et al. 2009). To quantify the nodal centrality for the graphs constructed from the rs-fMRI data, we used weighted DC (nodal strength). For a weighted graph, weighted DC is defined as the sum of the weights from edges connecting to a node. Thus, a node has a high degree if it is strongly connected to many other nodes.

Weighted DC was calculated at the individual level, and the voxel-wise maps were standardized to z-scores to allow comparisons across subjects and used as inputs of subsequent group-level analysis (hereby referred to as DC, if not otherwise indicated). Hence, these DC values are relative and reflect centrality relative to the rest of the brain, rather than absolute centrality. In order to account for differences in network cost influencing network topology (Achard and Bullmore 2007), network density (i.e the ratio between edges present in the network graph and the maximum number of edges that the graph can contain) was calculated for each subject to be included as a covariate in analyses. An average functional brain network was also constructed by converting each subject’s thresholded connectivity matrix into a Fisher Z matrix and averaging across participants. The averaged Fisher Z matrices were then back-transformed by using the inverse Fisher Z-transformation. DC was subsequently calculated on the average matrix. Using the same approach, weekly average matrices were also constructed from 38-41 weeks PMA. To ensure consistency across weeks, the same number of subjects were selected for each week (15 infants), with the lowest number of postnatal days as the selection criteria.

### Follow-up Outcome Data

Neurodevelopmental follow-up assessments were offered to all participants at around 18 months corrected age as part of the dHCP. Neurodevelopment was assessed with a number of behavioural and observational assessments, including the Bayley Scale of Infant and Toddler Development, Third Edition (Bayley 2006), The Child Behavior Checklist 1^1/2^-5 (CBCL; (Achenbach and Rescorla 2000) and The Quantitative Checklist for Autism in Toddlers (Q-CHAT; (Allison et al. 2008). The cognitive, motor and language composite scores of the Bayley Scale; the Externalising, Internalising and Other Behavioural Problems raw scores of the CBCL, and the total QCHAT score were included as outcome measures in this study. Index of multiple deprivation (IMD) was included as a covariate in statistical analyses. The IMD score is a composite score of social risk calculated based on the mother’s address at the time of birth (Abel et al. 2016). As not all infants completed all follow up assessments, the number of infants with data available for each measure is indicated in Table 1. Assessments were performed at St Thomas’ Hospital, London by two experienced assessors (authors AC, paediatrician and SF, chartered psychologist).

### Statistical Analyses

In the term infant group, the effect of PMA was explored in a general linear model (GLM) including postnatal days, sex, network density, total number of FD outliers and voxel-wise temporal signal to noise ratio (tSNR) as covariates. A groupwise comparison was conducted between preterm and term infants in a GLM including PMA at scan, sex, network density, total number of FD outliers and tSNR as covariates. The effect of GA at birth as a continuous variable was also explored in a GLM with the same covariates as in the groupwise analysis. A GLM assessing the interaction between group (term versus preterm) and PMA was also run to assess whether the effect of PMA at scan varies as a function of preterm birth. Network density and tSNR were included as covariates because subjects have varying network density (0.58±0.15 (mean±SD)), and the within-subject distribution of tSNR in the neonatal fMRI scans is not uniformly distributed. FD outliers were included as a covariate as head motion during fMRI data acquisition is known to cause signal artefact and affect estimates of functional connectivity (Satterthwaite et al. 2013). A version of the standard anatomical automatic labelling atlas (Tzourio-Mazoyer et al. 2002) adapted to the neonatal brain (Shi et al. 2011) was used for localization of results. Statistical analyses were carried out using nonparametric permutation inference as implemented in the FSL *randomise* tool, with 10000 permutations per test (Winkler et al. 2014). Threshold free cluster enhancement and family wise error rate was applied to correct for voxel-wise multiple comparisons (Smith and Nichols 2009).

The effects of PMA at scan and GA at birth were also explored within functional resting state networks (RSNs) previously identified in the neonatal brain (Eyre et al. 2021). Within the RSNs identified by Eyre et al. (2021), median DC (z-score) was extracted, and partial correlations were conducted between median DC and PMA at scan (term-born neonates, correcting for GA at birth, sex, total number of FD outliers and network density). In a multivariate model, the relationship between birth status (preterm versus term-born neonates) and median DC in the RSNs was assessed with PMA at scan, sex, FD outliers and network density as covariates. Partial correlations were also carried out between median DC and GA at birth as a continuous variable (preterm & term-born neonates combined, correcting for PMA at scan, sex, FD outliers and network density). Reported p-values were uncorrected, but false discovery rate (FDR) procedure (Benjamini and Hochberg 1995) was applied to highlight p-values surviving multiple comparisons correction.

The relationship between relative DC and outcome at 18 months was assessed both at the voxel- and RSN level. The relationships between voxel-wise DC and each outcome measure (Bayley-III subscales, CBCL subscales and total QCHAT (all corrected for age at follow up assessment)) were explored in GLMs including PMA at scan, postnatal days, sex, network density, total number of FD outliers, tSNR and IMD as covariates in the combined cohort of preterm- and term-born infants. To explore the relationship between DC and outcome at the RSN level, univariable analyses were conducted with partial correlations between median DC for each RSN and each outcome measure, correcting for PMA at scan, sex, network density, total number of FD outliers, IMD and age at follow up assessment as covariates. Multiple linear regression was also used to test if median DC in the 11 RSNs significantly predicted outcome, again correcting for PMA at scan, sex, network density, total number of FD outliers and IMD in the model. To assess whether the relationship between median DC and outcome differs with birth status (preterm versus term), univariate analyses were also conducted in the term-born cohort and the preterm-born cohort separately, both at the voxel- and RSN level, and the interaction between median DC in each RSN and birth status was included in a multivariable model.

### Data Availability

The dHCP is an open-access project. The imaging and collateral data used in this study were included in the 2021 (third) dHCP data release, which can be downloaded by completing a data usage agreement and registering at http://data.developingconnectome.org.

## Results

### Functional Centrality Distribution in Term-born Neonates

Analysis of the average brain connectivity matrices from term-born infants showed that the highest raw DC is evident within the left and right supramarginal gyrus, the left and right precentral gyrus and bilaterally in the cuneus, and lowest in the inferior temporal regions (Figure 1A). Panel B and C in Figure 1 show the weekly average DC maps as raw values and z-scored values, respectively.

**Figure 1.**
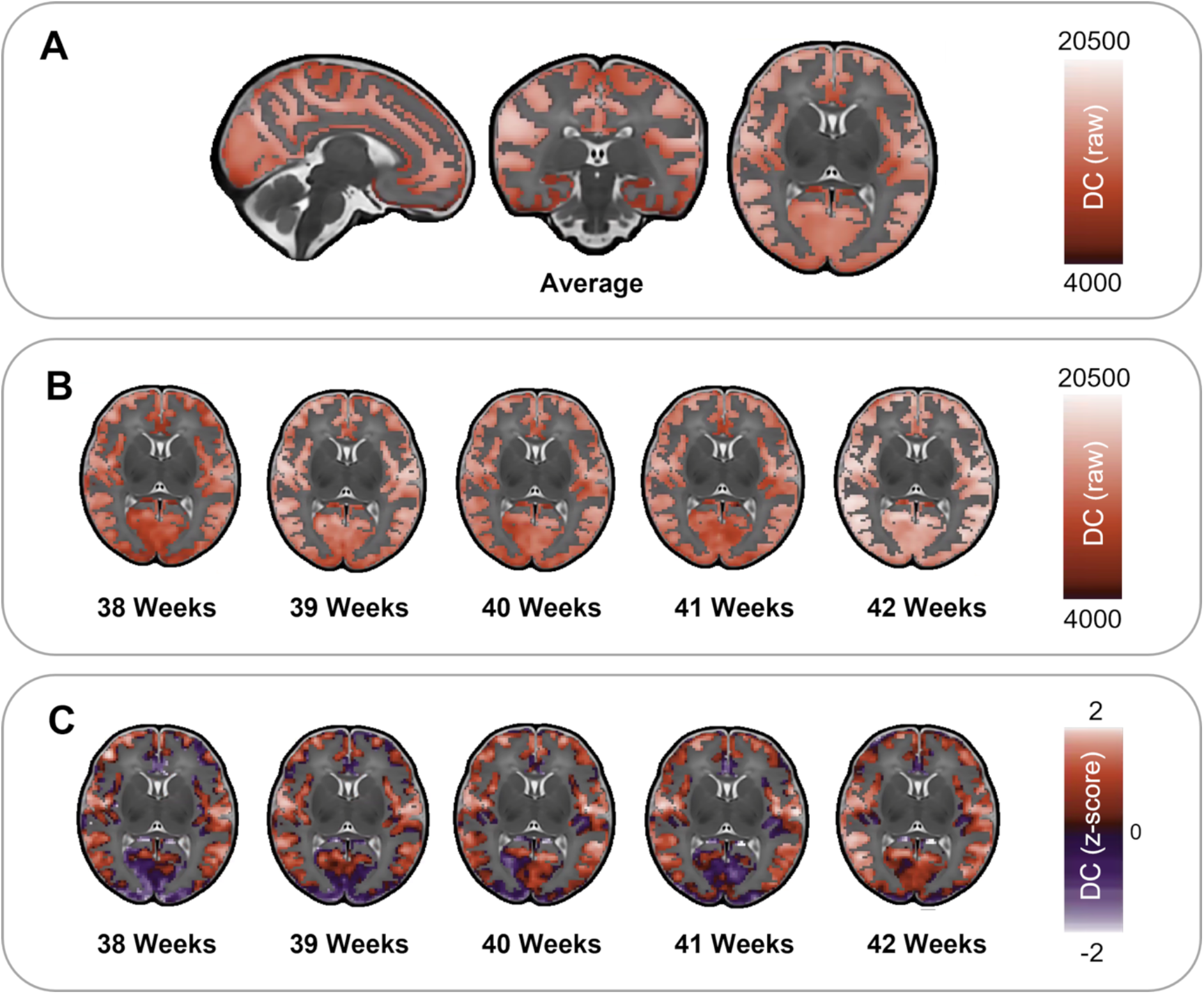
Average Degree distribution in term-born neonates (**A**, raw values) and average degree distribution across postnatal weeks (38-42 weeks, **B** = raw values, **C** = z-scored values).

### Association of functional centrality with age in Term-Born Neonates

In term-born infants, increasing PMA at scan was associated with an increase in voxelwise relative DC within the cuneus and calcarine fissure, and small regions within the lingual gyrus and superior occipital gyrus. PMA at scan was also associated with a decrease in relative DC in the bilateral precentral and postcentral gyrus, in addition to the rolandic operculum, insula and small regions within the transverse temporal gyrus (corrected for postnatal days, sex, FD motion outliers, network density and tSNR, Figure 2A).

**Figure 2.**
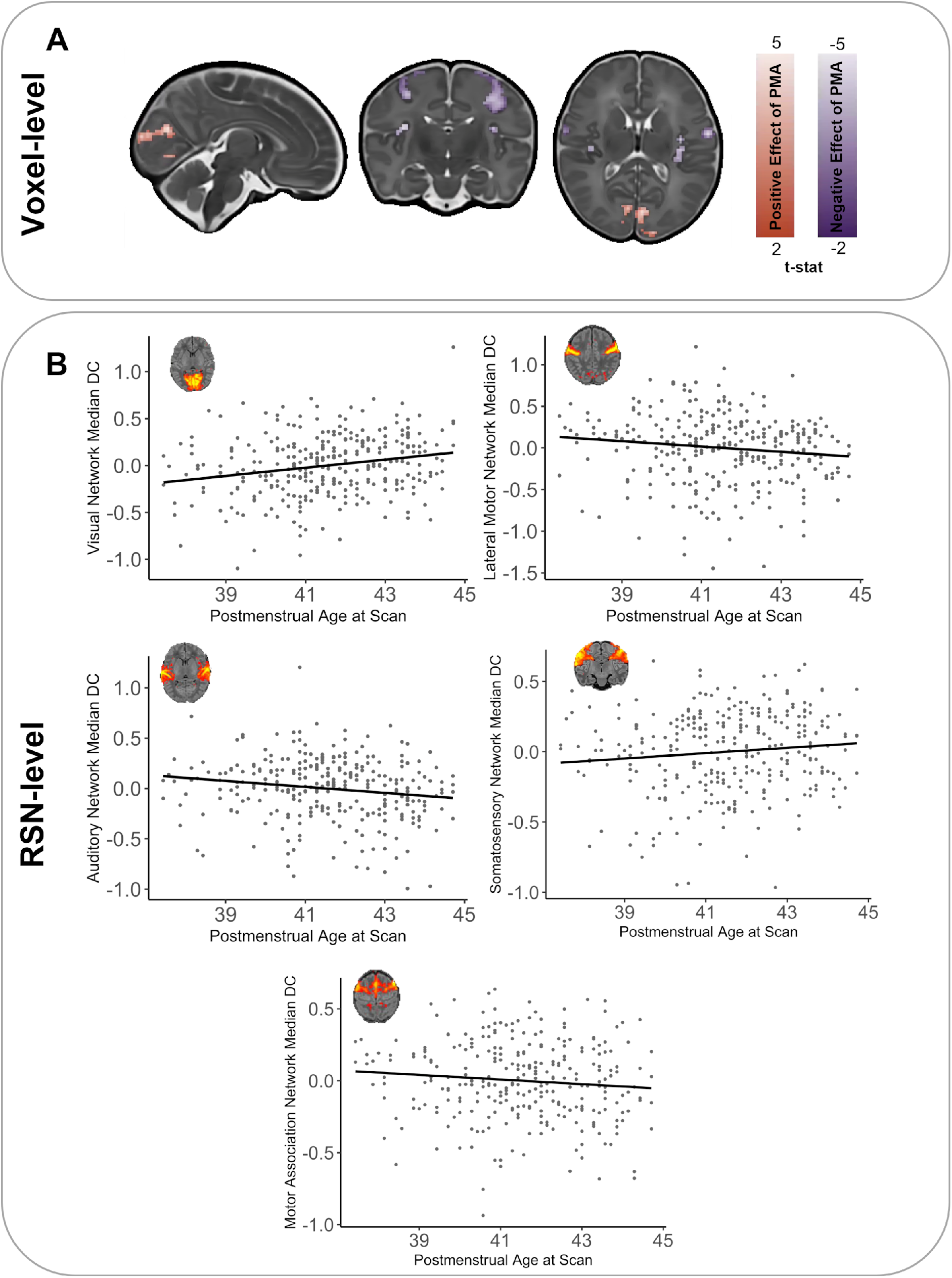
Effect of postmenstrual age (PMA) at scan on degree centrality in term born infants. **A)** PMA was associated with voxel-wise regional increases (red) and decreases (blue) in relative degree (T-statistic, p<0.025). **B)** Significant relationship were also present between the relative median degree centrality at the RSN-level and PMA (Z-score, residuals corrected for GA at birth, sex, FD outliers & density). RSN figures adapted from Eyre et.al., (2021).

A similar pattern of results was observed when exploring median DC in 11 established neonatal functional RSNs (Eyre et.al., 2021). After correcting for multiple comparisons, there was a significant positive relationship between the median relative DC and PMA in the visual RSN (r=0.30, p<0.001). A negative relationship was also observed in the auditory RSN (r=-0.22, p<0.001) and the lateral motor RSN (r=-0.17, p=0.003, Figure 2B). Correlations were also present within the somatosensory RSN and the motor association RSN (r=0.15, p=0.012 and r=-0.13, p=0.021, respectively).

### Comparing Functional Centrality in term and preterm infants

At the voxel-level, preterm-born infants showed higher relative DC in the right and left cuneus and the left and right calcarine fissure. Higher DC was also observed in very limited regions within the middle frontal gyrus and the inferior frontal gyrus (bilateral triangular part, right orbital part). Preterm-born infants showed lower centrality predominantly within the right cingulum and precunueus, in addition to the right rolandic operculum, the left paracentral lobule and the right and left precentral gyrus (corrected for PMA at scan, sex, motion outliers, network density and tSNR, Figure 3A).

**Figure 3.**
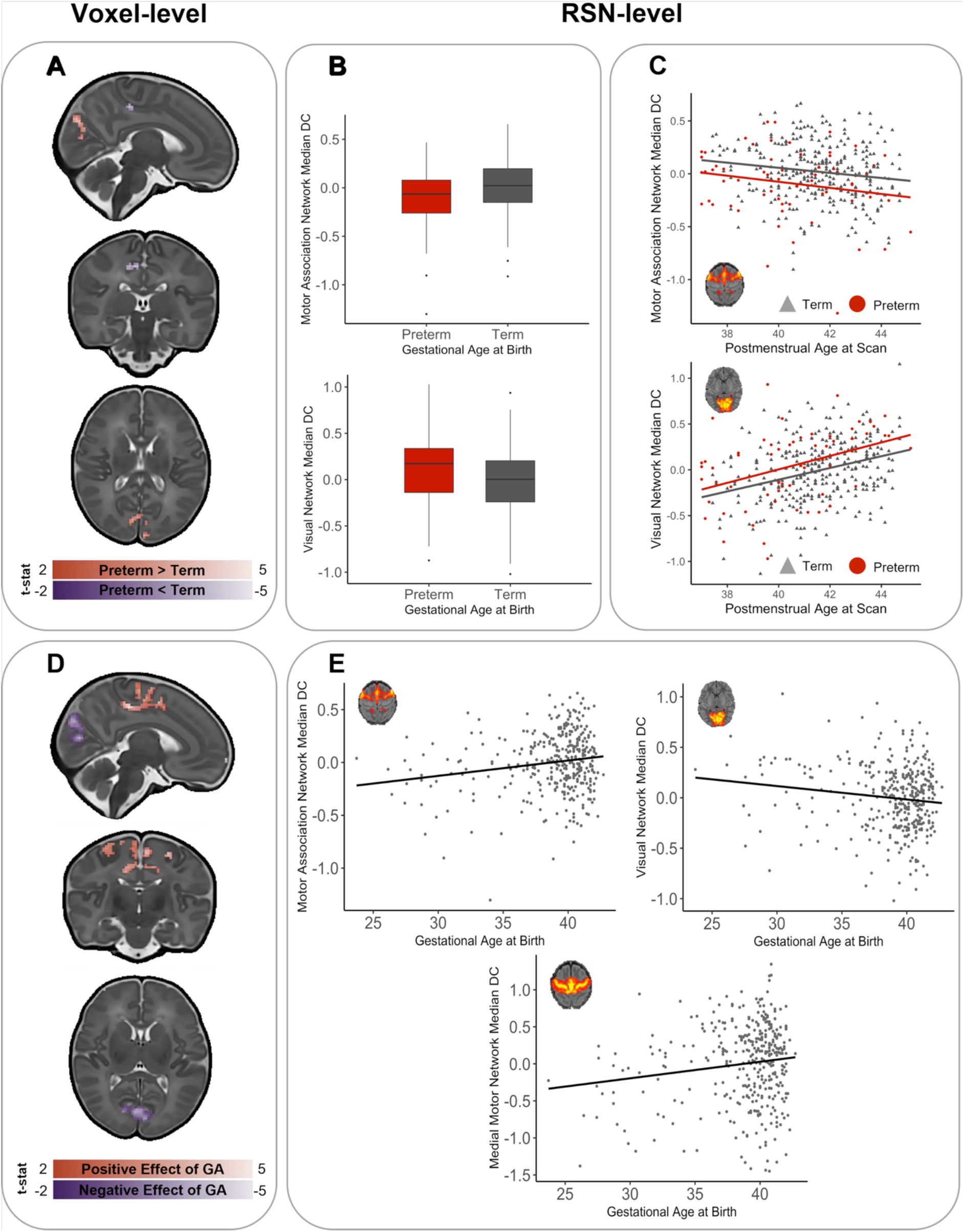
Effect of Preterm birth on degree centrality at TEA. **A)** Preterm-born infants show areas of voxel-wise regionally higher (red) and lower (blue) relative functional centrality compared to term-born infants**. B)** Significant differences were also present between preterm-born and term-born infants at the RSN level. **C)** There was no interaction between birth status (preterm-birth versus term-birth) and PMA at scan. **D)** When GA was treated as a continuous variable, GA at birth was associated with voxel-wise regional increases (red) and decreases (blue) in relative functional centrality (T-statistic, p<0.025) at term-equivalent age. **E)** Significant relationship was also present between the relative median degree centrality at the RSN-level and GA (Z=score, residuals, corrected for PMA at scan, sex, FD outliers and density. Representative RSN figures adapted from Eyre et.al., (2021).

When assessing median degree within established neonatal functional RSNs, preterm-born infants showed a decreased median DC in the motor association RSN (p<0.001) and increased centrality in the visual RSN (p=0.009) when compared with term-born controls (Figure 3B). There was no statistically significant interaction between group (term/preterm) and PMA at a voxel- or RSN level (Figure 3C).

In the combined cohort of term-born and preterm-born neonates at term-equivalent age, younger GA at birth resulted in lower voxel-wise relative DC in the bilateral supplementary motor areas, the bilateral precentral gyrus, the paracentral lobules and the right and left cingulum, in addition to the right precuneus. Younger GA at birth was also associated with increased relative centrality in the right and left calcarine fissure and cuneus, in addition to the left superior occipital gyrus, and very limited regions within the inferior frontal gyrus (corrected for PMA at scan, sex, motion outliers, network density and tSNR, Figure 3D).

At the RSN level, the median relative DC in the motor association RSN and medial motor RSN were positively related to GA at birth (r=0.19, p<0.001 and r=0.15, p=0.004, respectively), and median DC in the visual RSN was negatively associated with GA at birth (r=-0.15, p=0.004), significant after multiple comparisons correction (Figure 3E).

### Effect of network density and tSNR

To understand the effect of network density and tSNR on neonatal DC, the effect of including these variables in analysis was explored. Inclusion of network density and tSNR as covariates in the voxel-wise analyses did not significantly alter the location of results but did impact on the magnitude (see Supplementary Figures 1-3).

### The Relationship between DC and Outcome at 18 Months

There were no voxel-wise relationships between relative DC and neurodevelopmental outcomes at 18 months significant after multiple comparison correction in the term- and preterm-born infants combined, in the term-born infants only, or the preterm-born infants only. There were also no relationships that survived multiple comparison correction between DC and outcome at 18 months at the RSN level. Importantly, this was also the case for the regions affected by preterm birth at term equivalent age.

Prior to multiple comparison correction, at a significance threshold of p<0.05, some significant relationships were present. As this may be informative for future studies, these results have been included in the supplementary material. Briefly, prior to multiple comparison correction in the univariate analysis, CBCL Externalising was related to median DC in the visual association network (p=0.009, uncorrected), and CBCL *other* was associated with median DC in the auditory network (p=0.038, uncorrected) in the term- and preterm-born infants combined (Supplementary Table 1). In the term-born infants, Bayley Communication was associated with median DC in the posterior-parietal RSN (p=0.038, uncorrected), and CBCL Internalising was associated with median DC in the somatosensory RSN (p=0.039, uncorrected, Supplementary Table 2). In the preterm-born infants, CBCL Internalising was associated with median DC in the auditory RSN (p=0.018, uncorrected), Bayley Motor was associated with median DC in the prefrontal RSN (p=0.026, uncorrected), and CBCL Externalising was associated with the Visual Association RSN (p=0.049, uncorrected, Supplementary Table 3).

In the multivariable models, the only significant predictor prior to multiple comparison correction was the visual association RSN in the CBCL Externalising model (Supplementary Table 4). When the interaction between birth status (preterm versus term) and each of the 11 RSN networks were included in the models, significant interactions were present between median DC in the lateral motor RSN and birth status (preterm versus term), as well as the motor association RSN and birth status, when predicting CBCL externalising (Supplementary Table 5).

## Discussion

The aim of this study was to characterize postnatal functional network centrality development in term born infants at the voxel-level, in addition to exploring how centrality may be altered by preterm birth. Thus, we here provide a description of a voxel-wise measure of functional centrality (DC) in the neonatal brain and explore how it is associated with age at scan in term-born infants and GA at birth in a cohort of term-born and preterm-born infants. In the early postnatal period of term-born infants, our results suggest that the typical maturational trajectory involves an increase in relative functional centrality predominantly within the visual cortex, and a decrease in relative functional centrality within the primary motor and auditory cortices. Further, preterm birth appears to affect centrality largely in the same brain areas, although alterations in functional centrality associated with preterm birth in the motor cortex correspond to largely medial and association RSNs, compared to the lateral motor RSN being associated with age at scan in term infants. Comparing median values corresponding to previously defined neonatal RSNs (Eyre et al. 2021), preterm-born infants showed an increased functional centrality in the visual RSN and decreased centrality in the motor RSN. No statistically significant relationship was observed between functional centrality and developmental outcomes at 18 months.

The changes in relative functional centrality associated with age at scan in term-born infants are located within primary RSNs. The maturation of RSNs shows a primary-to-higher-order ontogenetic sequence, with RSNs encompassing primary motor, visual and auditory cortices already showing adult-like topology in term-born infants (Eyre et al. 2021). This is in line with neonates demonstrating visual processing capabilities albeit somewhat limited (Brémond-Gignac et al. 2011), and the capability for movement (Einspieler et al. 2008), and tactile and auditory processing (Maitre et al. 2020)(Ferronato et al. 2014) perinatally. Thus, the main changes in relative functional centrality associated with age at scan are within the RSNs which are the most substantially developed in early life and are important for early interactions with the environment.

Whilst spontaneous retinal activity allows the visual pathways to start to function early in the third trimester, following birth there is a large increase in visual stimuli, inducing increased cerebral activity and maturational mechanisms (Dubois et al. 2020). The fact that we observe an increase in relative centrality within the visual system may reflect the further improvements in visual acuity (Norcia and Tyler 1985; Hamer et al. 1989), contrast sensitivity (Pirchio et al. 1978), orientation selectivity (Morrone and Burr 1986) and motion sensitivity (Hadad et al. 2015) taking place over the first months following birth through establishment and maturation of the visual pathways (Siu and Murphy 2018). This is in line with previous seed-based connectivity studies suggesting a quantitative increase in connectivity of the visual RSN during the first months after birth (Gao et al. 2015).

Similar to the visual network, the primary motor and auditory networks show adult-like topology at birth (Keunen et al. 2017). In contrast to the visual RSN however, we observed a decrease in relative functional centrality associated with age at scan within these RSNs. This may reflect an experience-expectant process whereby broad cortical circuits become increasingly refined with exposure, resulting in postnatal specialisation of connectivity. However, it is worth noting that previous work looking specifically at speech processing (Shultz et al. 2014) and auditory stimuli more generally (Gao et al. 2015) suggests that this process takes place slightly later than the age of infants included in the present study, warranting a more fine-grained assessment of the temporal trajectories of specialization within these RSNs to assess whether the changes we observe reflect refinement or a relative increase in the importance of other RSNs.

With the brain undergoing dramatic changes in functional activity during the third trimester, infants born prematurely during this sensitive period are vulnerable to the influence of developmentally unexpected sensory stimuli which is not age-appropriate and significantly different from the womb in duration, complexity and intensity (Beltrán et al. 2021; Mellado et al. 2021). Such exposure to inadequate and/or inappropriate visual, auditory and motor stimuli may contribute to subtle brain changes both in primary cortices and associative areas, which can in turn translate into long-lasting sensorimotor alterations (El-Metwally and Medina 2020). Indeed, children who are born prematurely more often present with conditions related to disrupted sensory processing, such as ADHD and autism and learning and memory problems (El-Metwally and Medina 2020) in addition to neuromotor abnormalities (Wheelock et al. 2018).

Notably, the areas that appear most affected by preterm birth in the present study are also predominantly located within the motor and visual RSNs. This suggests that preterm birth affects the regions which are showing the most substantial development during the perinatal period. Given the crucial role of sensory input in neurodevelopment (Mellado et al. 2021), preterm infants exposed to developmentally unexpected stimulation of motor and visual networks (i.e *ex utero* versus *in utero* experience) may show accelerated maturation along normal developmental trajectories or adopt adverse and/or compensatory mechanisms as an adaptive response to cope with the environment (De Asis-Cruz et al. 2020; Eyre et al. 2021; Mellado et al. 2021). This may in turn result in the altered patterns of functional centrality we observe in preterm-born infants in the current study. However, it is challenging to disentangle whether the alterations in functional centrality observed in preterm-born infants are a result of less time spent *in utero* and the consequent premature exposure to the postnatal environment or underlying clinical and/or genetic factors associated with preterm-birth.

As the preterm-born infants included in this study are moderate or late preterm, it is noteworthy that differences in functional architecture are present even in this cohort. This evidences the importance of studying preterm-birth in a wide range of GAs at birth and not solely infants born very or extremely preterm, particularly as 80% of preterm infants are born moderate or late preterm (Blencowe et al. 2012). It is however also worth noting that the present study population is a cohort of “healthy” preterm-born infants, with no incidental findings of likely clinical significance, and who did not demonstrate significantly different developmental outcomes from term-born infants at 18 months. We are therefore unable to extrapolate our findings to more severely affected preterm babies, such as those with significant white matter damage.

The fact that we did not observe any robust relationships between functional centrality and outcome at 18 months that survived multiple comparison correction is in line with the majority of neonatal fMRI studies, where clear relationships between functional connectivity patterns in the perinatal period and outcome at 1-2 years of age are seldom reported, particularly in term-born cohorts. Detecting links between brain imaging and general measures of behaviour is difficult in the developing brain, as the changes associated with normal maturation are much larger than those associated with individual differences (Batalle et al. 2018). The functional differences between individuals may be so subtle that they do not manifest behaviourally this early on, or the relationships may be more complex and non-linear. There is also the possibility that subtle differences may be influenced or magnified by on-going environmental exposures throughout development (Livingston and Happé 2017) and that small functional differences in primary networks could disrupt or unbalance the developmental trajectories of higher order networks (Ciarrusta et al. 2020) leading to behavioural manifestations later in development. Additionally, it is conceivable that the relationship between functional alterations associated with preterm birth and outcome might have been clearer in a cohort of extremely preterm infants and/or infants with more severe damage as a result of preterm birth. Indeed, previous studies have shown relationships between functional connectivity at term equivalent age and outcome at 1-2 years of age in very preterm infants (Toulmin et al. 2021). It may also be the case that disruption to functional centrality topology observed at term-equivalent age in preterm-born infants may be ‘normalised’ or be compensated for over the first year(s) of life, so that early disruption does not result in behavioural consequences at 18 months.

Our study advances on previous work by providing a description of functional centrality in a large, representative sample. Our results offer complementary information to the findings by Cao et al. (2017), who demonstrated increases in absolute DC predominantly within the precentral and postcentral gyri and supplementary motor area and decreases within the posterior cingulate and precuneus cortex from 31 to 42 weeks PMA. The current findings provide a picture of the alterations in relative centrality that are associated with preterm birth at term-equivalent age. Although our results show somewhat different patterns of altered centrality associated with preterm birth than those reported by Gozdas et al. (2018), a direct comparison is not possible due to differences in methodology and sample - most notably, their comparison between groups being carried out on a regional basis using 90 ROIs rather than a voxel-based approach.

Our study has several strengths. We use a large cohort and employ a metric that does not rely on a particular parcellation template or selection of a priori regions of interest. Additionally, whilst acknowledging that the role of network density in graph theory-based analyses is a complex issue (Hallquist and Hillary 2019), we are able to make some inferences about the importance of network density in our analyses. The inclusion of density as a covariate did have an effect on magnitude of our findings, but did not change their location, suggesting that the observed regional differences are not solely due to differences in global network topology.

The imaging system for the dHCP is also optimized for neonatal imaging. The close-fitting custom head coil provides exceptional signal to noise ratio (SNR) on the cortical surface (Hughes et al. 2017). However, this does result in a bias towards surface-proximate sources, which is further compounded by the use of highly accelerated multiband EPI (Fitzgibbon et al. 2020). Because of this, we did not include deep grey matter in our analysis, including regions such as the amygdala, thalamus and cerebellum. Additionally, despite the dHCP functional pipeline including advanced distortion correction techniques (Fitzgibbon et al. 2020), some signal is lost due to air/tissue and bone/tissue interfaces. The use of a single phase-encode direction (anterior-posterior) may also compress signal in frontal regions. To mitigate these effects, voxel-wise tSNR was included as a covariate in voxel-wise analyses. As motion is known to have an impact on the rs-fMRI signal (Power et al. 2012; Satterthwaite et al. 2012) and the term-born infants presented with more motion than the preterm-born infants, we cannot exclude the possibility that motion artefacts may have had an impact on our results. However, the dHCP functional pipeline includes multiple steps to mitigate the effect of motion and physiological confounds to minimise data loss; in addition, we excluded babies with high motion and included number of FD outliers as a covariate in analyses. These steps should also have helped mitigate the effect of differences in sleep state and arousal as these manifest in different levels of subject motion (Horovitz et al., 2009). We are however unable to account for any fundamental differences in BOLD signal in term-born and preterm-born infants linked to cerebrovascular factors (Bouyssi-Kobar et al. 2018).

Our study is also limited by the fact that our outcome assessments are only available at 18 months of age, meaning we are unable to establish whether the patterns of atypical functional connectivity we observe to be associated with preterm birth translate into behavioural or socio-emotional difficulties commonly observed in preterm-born infants in later life, or whether they reflect a transient pattern of altered functional architecture. Further work is needed to understand whether DC at birth holds any predictive value for behavioural outcomes at later timepoints. Lastly, although DC is a widely used method to study functional network centrality, various other centrality measures exist and may capture other aspects of functional connectivity (Zuo et al. 2012).

In summary, this work contributes to our understanding of typical maturational trajectories of functional centrality at the voxel and RSN level and explores how functional centrality is affected by preterm birth. Our findings suggest that the changes in centrality associated with typical maturation and preterm birth are both predominantly located within primary motor and visual cortices. This suggests that preterm birth largely affects the pattern of functional centrality of the cortical regions that undergo the most substantial development in the perinatal period, highlighting both the sensitive and significant nature of these regions during early life. As functional centrality was not related to performance on outcome measures at 18 months, further work is needed to ascertain whether the alterations in functional centrality associated with preterm birth are adaptive and serve a compensatory role, or whether they are disruptive and potentially predictive of neurodevelopmental features that are more subtle than those measured here at 18 months, or that emerge later in life.

## Supplementary Material

**Supplementary Figure 1.**
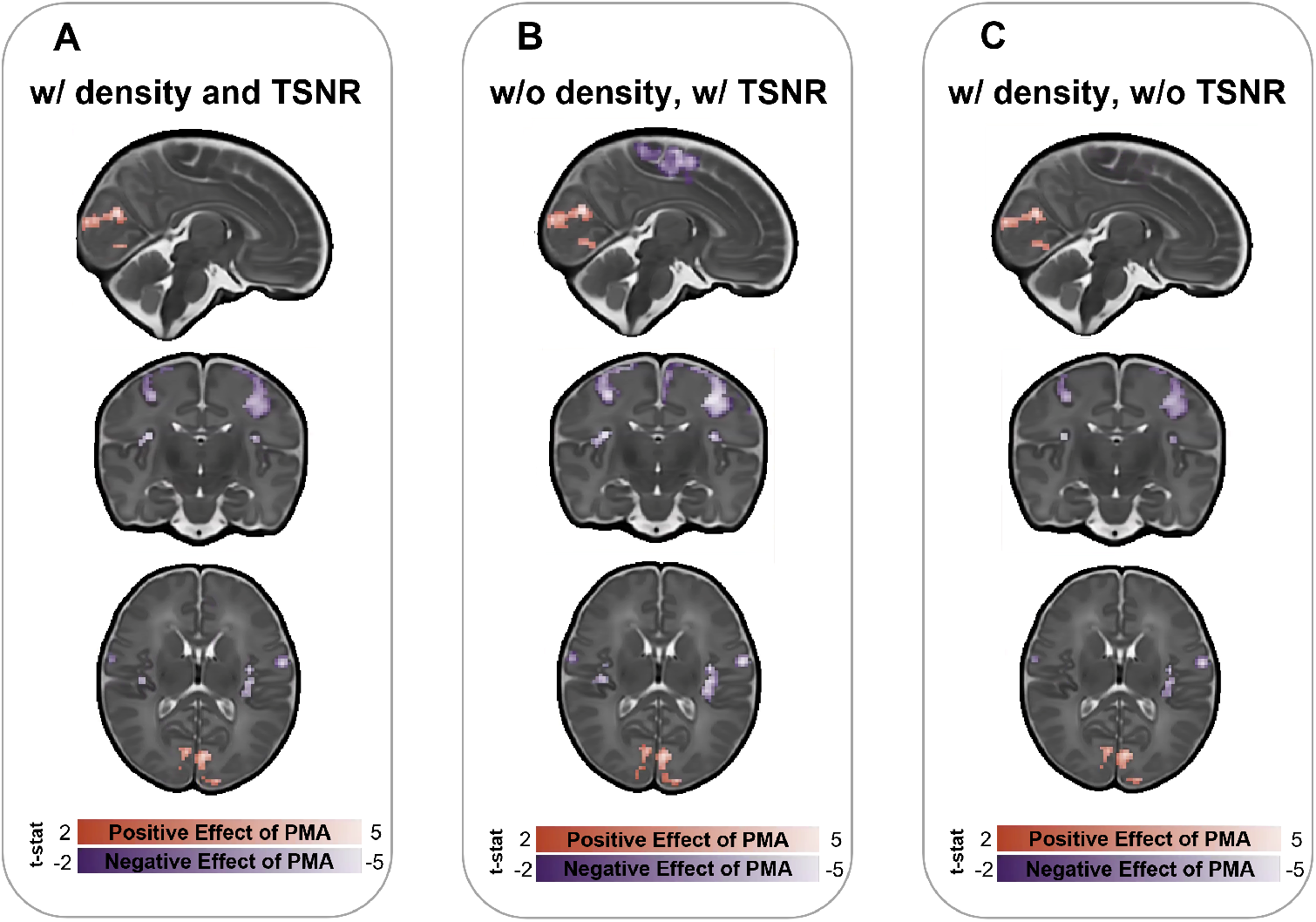
Postmenstrual age at scan, including density and tSNR as covariates **(A),** including tSNR but not density **(B),** and including density but not tSNR **(C).**

**Supplementary Figure 2.**
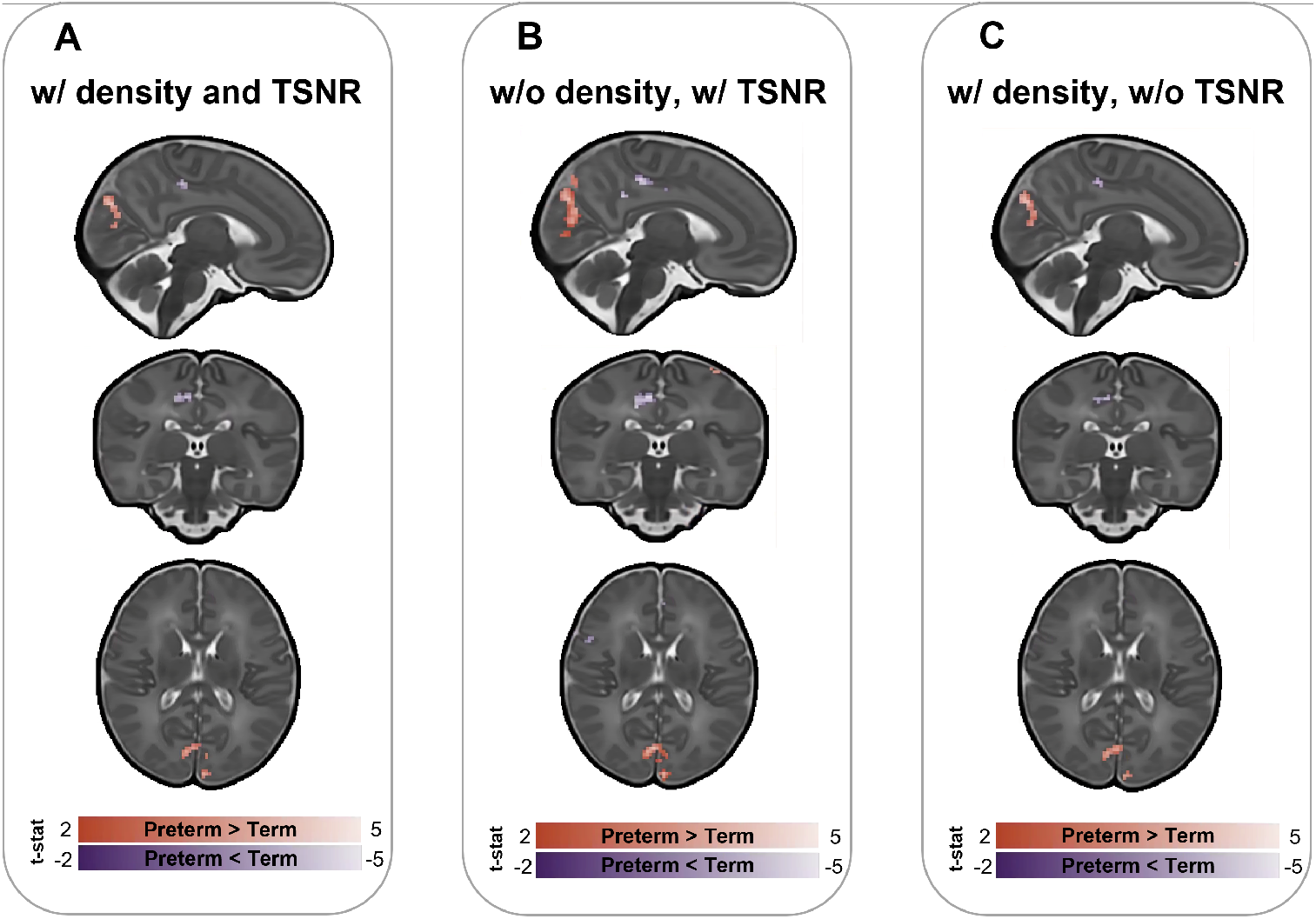
Preterm versus term, including density and tSNR as covariates **(A),** including tSNR but not density **(B),** and including density but not tSNR **(C).**

**Supplementary Figure 3.**
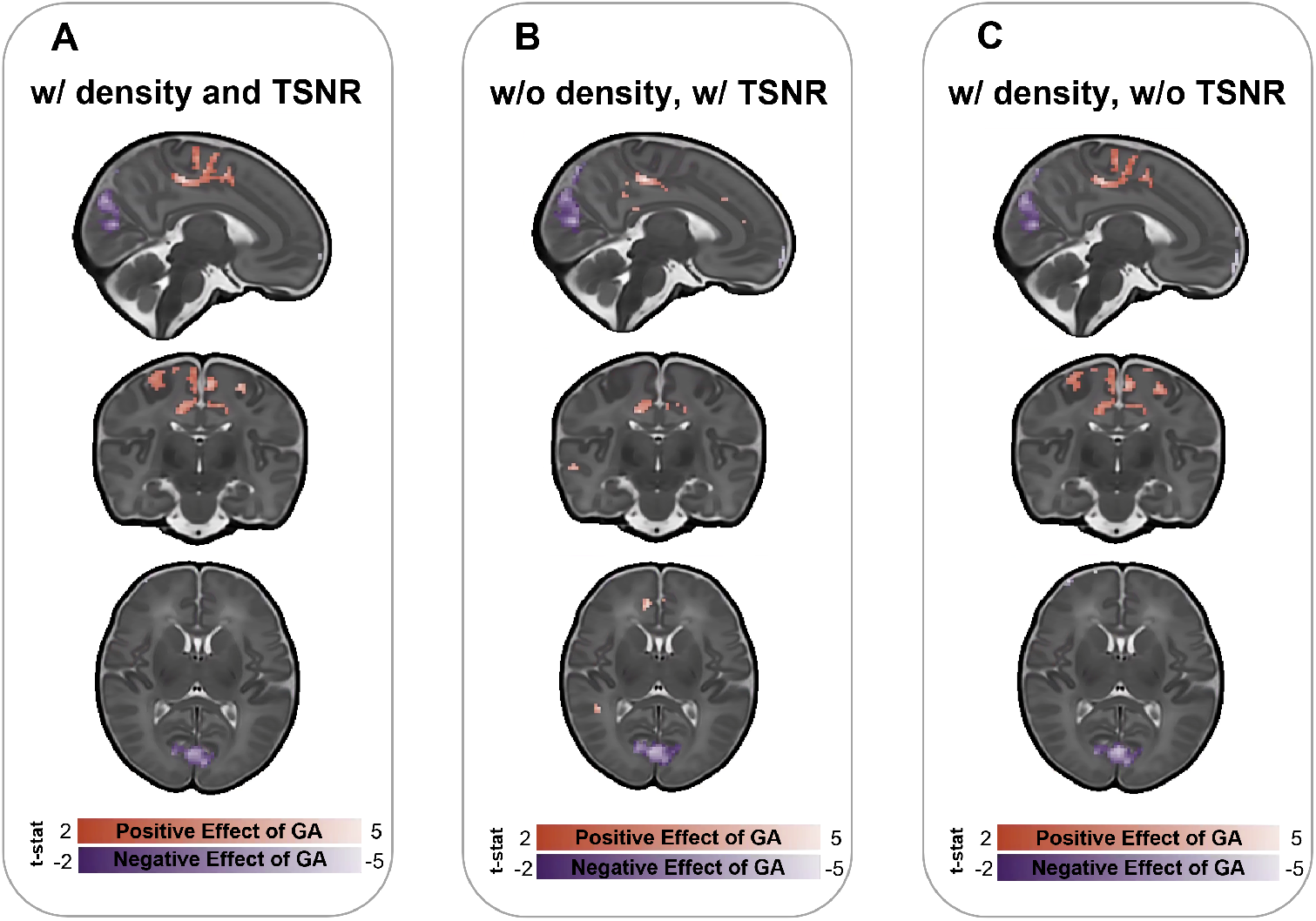
Gestational age at birth, including density and tSNR as covariates (**A)**, including tSNR but not density **(B)**, and including density but not tSNR **(C).**

**Supplementary Table 1.**
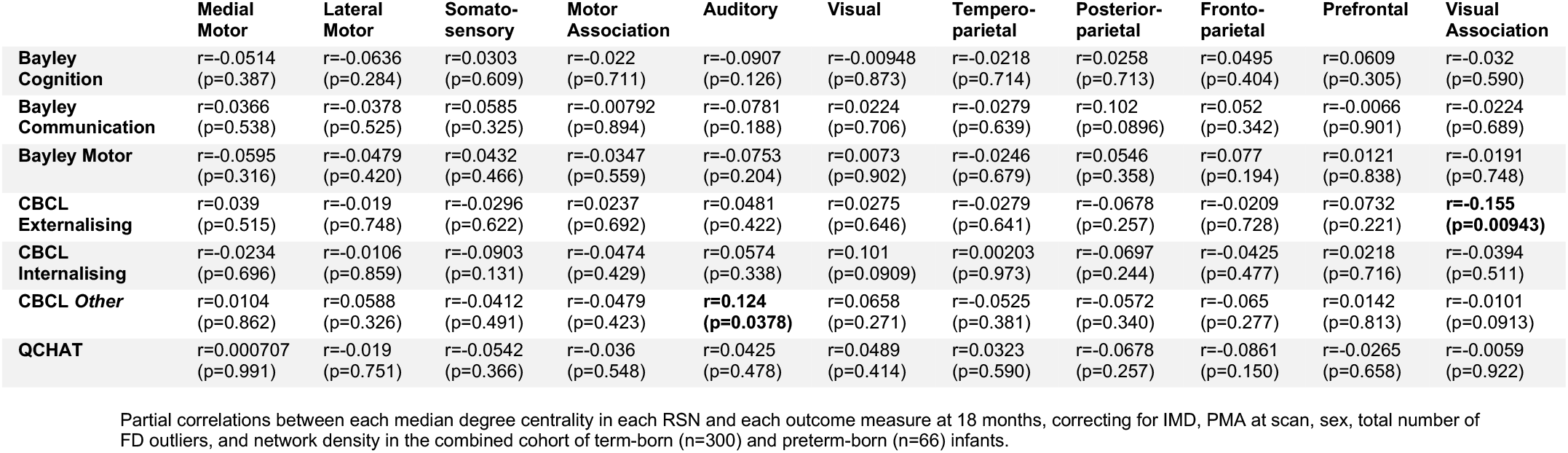
Univariable Analyses – term & preterm.

|  | Medial Motor | Lateral Motor | Somato-sensory | Motor Association | Auditory | Visual | Temporo-parietal | Posterior-parietal | Fronto-parietal | Prefrontal | Visual Association |
| --- | --- | --- | --- | --- | --- | --- | --- | --- | --- | --- | --- |
| Bayley Cognition | r=-0.0514<br>(p=0.387) | r=-0.0636<br>(p=0.284) | r=0.0303<br>(p=0.609) | r=-0.022<br>(p=0.711) | r=-0.0907<br>(p=0.126) | r=-0.00948<br>(p=0.873) | r=-0.0218<br>(p=0.714) | r=0.0258<br>(p=0.713) | r=0.0495<br>(p=0.404) | r=0.0609<br>(p=0.305) | r=-0.032<br>(p=0.590) |
| Bayley Communication | r=0.0366<br>(p=0.538) | r=-0.0378<br>(p=0.525) | r=0.0585<br>(p=0.325) | r=-0.00792<br>(p=0.894) | r=-0.0781<br>(p=0.188) | r=0.0224<br>(p=0.706) | r=-0.0279<br>(p=0.639) | r=0.102<br>(p=0.0896) | r=0.052<br>(p=0.342) | r=-0.0066<br>(p=0.901) | r=-0.0224<br>(p=0.689) |
| Bayley Motor | r=-0.0595<br>(p=0.316) | r=-0.0479<br>(p=0.420) | r=0.0432<br>(p=0.466) | r=-0.0347<br>(p=0.559) | r=-0.0753<br>(p=0.204) | r=0.0073<br>(p=0.902) | r=-0.0246<br>(p=0.679) | r=0.0546<br>(p=0.358) | r=0.077<br>(p=0.194) | r=0.0121<br>(p=0.838) | r=-0.0191<br>(p=0.748) |
| CBCL Externalising | r=0.039<br>(p=0.515) | r=-0.019<br>(p=0.748) | r=-0.0296<br>(p=0.622) | r=0.0237<br>(p=0.692) | r=0.0481<br>(p=0.422) | r=0.0275<br>(p=0.646) | r=-0.0279<br>(p=0.641) | r=-0.0678<br>(p=0.257) | r=-0.0209<br>(p=0.728) | r=0.0732<br>(p=0.221) | **r=-0.155**<br>**(p=0.00943)** |
| CBCL Internalising | r=-0.0234<br>(p=0.696) | r=-0.0106<br>(p=0.859) | r=-0.0903<br>(p=0.131) | r=-0.0474<br>(p=0.429) | r=0.0574<br>(p=0.338) | r=0.101<br>(p=0.0909) | r=0.00203<br>(p=0.973) | r=-0.0697<br>(p=0.244) | r=-0.0425<br>(p=0.477) | r=0.0218<br>(p=0.716) | r=-0.0394<br>(p=0.511) |
| CBCL Other | r=0.0104<br>(p=0.862) | r=0.0588<br>(p=0.326) | r=-0.0412<br>(p=0.491) | r=-0.0479<br>(p=0.423) | **r=0.124**<br>**(p=0.0378)** | r=0.0658<br>(p=0.271) | r=-0.0525<br>(p=0.381) | r=-0.0572<br>(p=0.340) | r=-0.065<br>(p=0.277) | r=0.0142<br>(p=0.813) | r=-0.0101<br>(p=0.0913) |
| QCHAT | r=0.000707<br>(p=0.991) | r=-0.019<br>(p=0.751) | r=-0.0542<br>(p=0.366) | r=-0.036<br>(p=0.548) | r=0.0425<br>(p=0.478) | r=0.0489<br>(p=0.414) | r=0.0323<br>(p=0.590) | r=-0.0678<br>(p=0.257) | r=-0.0861<br>(p=0.150) | r=-0.0265<br>(p=0.658) | r=-0.0059<br>(p=0.922) |

Partial correlations between each median degree centrality in each RSN and each outcome measure at 18 months, correcting for IMD, PMA at scan, sex, total number of FD outliers, and network density in the combined cohort of term-born (n=300) and preterm-born (n=66) infants.

**Supplementary Table 2.**
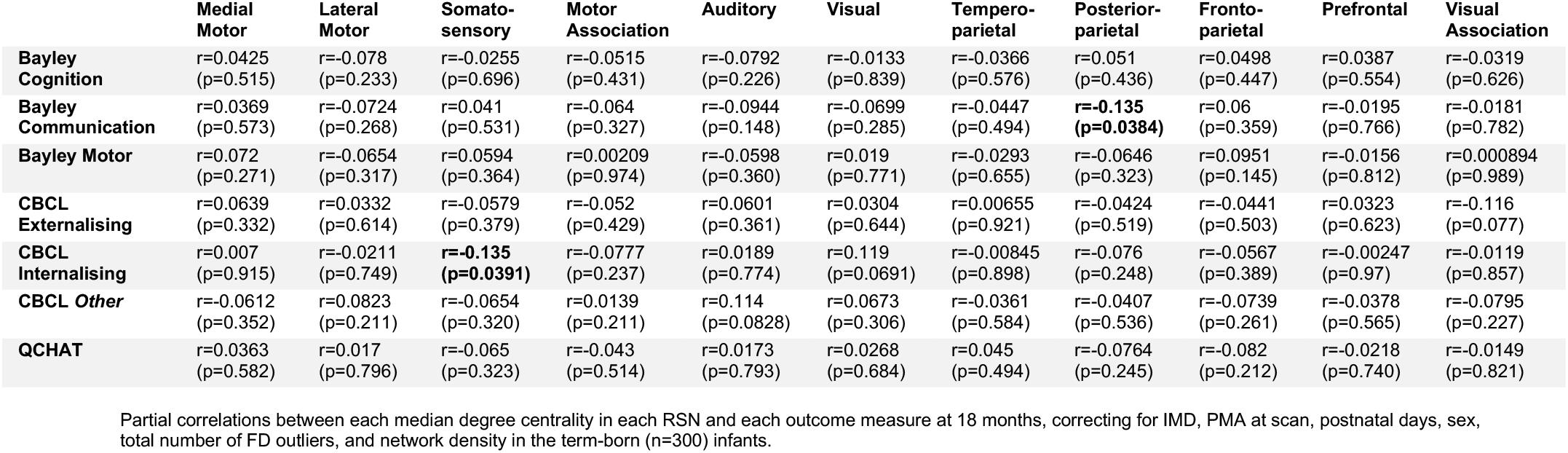
Univariable Analyses – term only.

**Supplementary Table 3.** Univariable Analyses – preterm only.

Univariable Analyses –preterm only
|  | Medial Motor | Lateral Motor | Somato-sensory | Motor Association | Auditory | Visual | Tempero-parietal | Posterior-parietal | Fronto-parietal | Prefrontal | Visual Association |
| --- | --- | --- | --- | --- | --- | --- | --- | --- | --- | --- | --- |
| Bayley Cognition | r=0.0629<br>(p=0.681) | r=-0.0818<br>(p=0.593) | r=-0.00019<br>(p=0.999) | r=0.115<br>(p=0.453) | r=-0.148<br>(p=0.333) | r=-0.198<br>(p=0.191) | r=-0.0398<br>(p=0.795) | r=-0.158<br>(p=0.301) | r=-0.0695<br>(p=0.650) | r=0.246<br>(p=0.104) | r=-0.0562<br>(p=0.714) |
| Bayley Communication | r=-0.168<br>(p=0.277) | r=0.0731<br>(p=0.637) | r=0.0488<br>(p=0.753) | r=0.147<br>(p=0.343) | r=-0.112<br>(p=0.470) | r=-0.118<br>(p=0.446) | r=-0.0366<br>(p=0.814) | r=-0.0283<br>(p=0.855) | r=0.0495<br>(p=0.749) | r=-0.229<br>(p=0.135) | r=0.0269<br>(p=0.863) |
| Bayley Motor | r=-0.181<br>(p=0.239) | r=-0.009<br>(p=0.954) | r=-0.0626<br>(p=0.686) | r=-0.198<br>(p=0.198) | r=-0.214<br>(p=0.163) | r=-0.0662<br>(p=0.669) | r=-0.113<br>(p=0.467) | r=-0.163<br>(p=0.291) | r=-0.114<br>(p=0.461) | **r=-0.335**<br>**(p=0.0264)** | r=-0.135<br>(p=0.383) |
| CBCL Externalising | r=-0.104<br>(p=0.506) | r=-0.290<br>(p=0.0589) | r=0.0153<br>(p=0.923) | r=0.272<br>(p=0.0778) | r=-0.0169<br>(p=0.914) | r=0.0159<br>(p=0.919) | r=-0.190<br>(p=0.222) | r=-0.194<br>(p=0.213) | r=-0.00382<br>(p=0.981) | r=0.242<br>(p=0.118) | **r=-0.303**<br>**(p=0.0485)** |
| CBCL Internalising | r=-0.106<br>(p=0.498) | r=0.0609<br>(p=0.698) | r=0.115<br>(p=0.463) | r=0.0625<br>(p=0.690) | **r=0.359**<br>**(p=0.0181)** | r=-0.0663<br>(p=0.673) | r=-0.00357<br>(p=0.982) | r=-0.180<br>(p=0.249) | r=-0.173<br>(p=0.266) | r=0.0834<br>(p=0.595) | r=-0.219<br>(p=0.159) |
| CBCL Other | r=-0.134<br>(p=0.393) | r=-0.115<br>(p=0.464) | r=0.0966<br>(p=0.538) | r=0.157<br>(p=0.315) | r=0.222<br>(p=0.153) | r=0.0366<br>(p=0.816) | r=-0.164<br>(p=0.292) | r=-0.222<br>(p=0.153) | r=-0.129<br>(p=0.411) | r=0.119<br>(p=0.447) | r=-0.224<br>(p=0.149) |
| QCHAT | r=-0.0965<br>(p=0.538) | r=-0.143<br>(p=0.361) | r=0.0173<br>(p=0.912) | r=-0.267<br>(p=0.083) | r=0.223<br>(p=0.151) | r=0.143<br>(p=0.361) | r=0.015<br>(p=0.924) | r=-0.0126<br>(p=0.936) | r=-0.0577<br>(p=0.713) | r=-0.0769<br>(p=0.624) | r=-0.0346<br>(p=0.826) |

Partial correlations between each median degree centrality in each RSN and each outcome measure at 18 months, correcting for IMD, PMA at scan, sex, total number of FD outliers, and network density in the preterm-born (n=66) infants.

**Supplementary Table 4.** Multivariable Analyses – w/o interaction terms.

Multivariable Analyses – w/o interaction terms
| Outcome Variable | Overall Model | Significant RSN predictors |
| --- | --- | --- |
| Bayley Cognition | F(17,273)=2.075, p=0.008, multiple R<sup>2</sup> = 0.1144, adjusted R<sup>2</sup> = 0.0593 |  |
| Bayley Communication | F(17,273)=1.906, p=0.018, multiple R<sup>2</sup> = 0.1061, adjusted R<sup>2</sup> = 0.0504 |  |
| Bayley Motor | F(17,273)=1.474, p=0.1035 (NS), multiple R<sup>2</sup> = 0.0841, adjusted R<sup>2</sup> = 0.0270 |  |
| CBCL Externalising | F(17,268)=1.725, p=0.039, multiple R<sup>2</sup> = 0.0986, adjusted R<sup>2</sup> = 0.0415 | Visual Association Network (b=-0.275, t=-2.562, p=0.011) |
| CBCL Internalising | F(17,268)=1.314, p=0.183 (NS), multiple R<sup>2</sup> = 0.077, adjusted R<sup>2</sup> = 0.0184 |  |
| CBCL Other | F(17,268)=1.284, p=0.202 (NS), multiple R<sup>2</sup> = 0.075, adjusted R<sup>2</sup> = 0.0167 |  |
| QCHAT | F(17,269)=1.353, p=0.160 (NS), multiple R<sup>2</sup> = 0.079, adjusted R<sup>2</sup> = 0.0206 |  |

Multiple linear regressions to predict each outcome measure, given the 11 RSNs, PMA at scan, sex, total number of FD outliers and network density in the combined cohort of term-born (n=300) and preterm-born (n=66) infants.

**Supplementary Table 5.** Multivariable Analyses – w/ interaction terms.

Multivariable Analyses – w/ interaction terms
| Outcome Variable | Overall Model | Significant RSN predictors |
| --- | --- | --- |
| Bayley Cognition | F(28,262)=1.147, p=0.2844 (NS), multiple R<sup>2</sup> = 0.1092, adjusted R<sup>2</sup> = 0.01396 |  |
| Bayley Communication | F(28,262)=1.378, p=0.1034 (NS), multiple R<sup>2</sup> = 0.1284, adjusted R<sup>2</sup> = 0.0352 |  |
| Bayley Motor | F(28,262)=0.9908, p=0.4829 (NS), multiple R<sup>2</sup> = 0.0957, adjusted R<sup>2</sup> = 0.0009 |  |
| CBCL Externalising | F(28,257)=1.742, p=0.01413, multiple R<sup>2</sup> = 0.1595, adjusted R<sup>2</sup> = 0.06797 | Lateral Motor (b=-0.5048, t=-2.510, p=0.01268), Motor Association (b=0.3302, t=-2.016, p=0.0448), Lateral Motor*Birth Status (b=0.4836, t=-2.198, p=0.02885), Motor Association*Birth Status (b=-0.5015, t=-2.899, p=0.00407) |
| CBCL Internalising | F(28,257)=1.192, p=0.2378 (NS), multiple R<sup>2</sup> = 0.115, adjusted R<sup>2</sup> = 0.01854 |  |
| CBCL Other | F(28,257)=1.290, p=0.1565 (NS), multiple R<sup>2</sup> = 0.1232, adjusted R<sup>2</sup> = 0.02771 |  |
| QCHAT | F(28,257)=1.488, p=0.05927 (NS), multiple R<sup>2</sup> = 0.1395, adjusted R<sup>2</sup> = 0.0457 |  |

Multiple linear regressions to predict each outcome measure, given the 11 RSNs, birth status, PMA at scan, sex, total number of FD outliers and network density in the combined cohort of term-born (n=300) and preterm-born (n=66) infants.

